# The indoor environment - a potential source for intact human-associated anaerobes

**DOI:** 10.1101/2020.12.02.406132

**Authors:** Manuela-Raluca Pausan, Marcus Blohs, Alexander Mahnert, Christine Moissl-Eichinger

**Author notes:** Corresponding author Correspondence to C. Moissl-Eichinger. Authors contributed equally (Manuela-Raluca Pausan, Marcus Blohs). Steigerwald Arzneimittelwerk GmbH, Bayer Consumer Health, Havelstrasse 5, D-64295 Darmstadt, Germany (Manuela-Raluca Pausan).

## Abstract

**Background:** People in westernised countries spend most of their time indoors. A healthy human microbiome relies on the interaction with and exchange of microbes that takes place between the human body and its environment. For this reason, the built environment might represent a potent source of commensal microbes. Anaerobic microbes are of particular interest, as researchers have not yet sufficiently clarified how the human microbiome acquires oxygen-sensitive microbes, such as obligate or facultative anaerobes.

**Methods:** We sampled ten households and used propidium monoazide to assess the viability of the collected prokaryotes. We compared the microbiome profiles based on 16S rRNA gene sequencing and confirmed our results by genetic and cultivation-based analyses.

**Results:** Quantitative and qualitative analysis revealed that most of the microbial taxa are of human origin. Less than 25% of the prokaryotic signatures found in built environment (BE) samples originate from intact – and thus potentially living – cells, indicating that aerobic and stress resistant taxa display an apparent survival advantage. Although the dominant microbial fraction identified on the bathroom floors is composed of aerobes, we confirmed the presence of strictly anaerobic taxa, including methanogenic archaea, in PMA-treated samples. As methanogens are regarded as highly sensitive to aerobic conditions, oxygen-tolerance experiments were performed with human-associated isolates to validate their survival. These results show that these taxa have a limited but substantial ability to survive in the BE. We determined that human-associated methanogens can survive oxic conditions for at least 6 h.

**Conclusions:** This study enabled us to collect strong evidence that supports the hypothesis that obligate anaerobic taxa can survive in the BE for a limited amount of time. This suggests that the BE serves as a potential source of anaerobic human commensals.

## Background

The development of a healthy human microbiome depends on extensive microbial transmission from other hosts, food, water, but also reservoirs in the built environment (BE; Browne et al., 2017). Especially for newborn infants, the acquisition of microbes is a crucial process, as they represent an empty niche for a variety of microorganisms. In this way, they differ from adults, who usually have well-established, complex and stable microbiomes. A fairly stable microbiome is normally achieved by the third year after birth [2] and, up until this point, numerous microorganisms are acquired from other microbiomes. Various factors affect this maturation process, including its speed and quality. These factors include the birth mode (Caesarean section vs. vaginal birth), type of feeding (breast-fed vs. formula-fed), medication (e.g. antibiotics), time point of introduction of solid food, quality of social interactions and contact to animals and nature [3–6]. During the maturation of the intestinal microbiome, facultative anaerobic microorganisms, which initially thrive in the gastrointestinal tract (GIT) after birth, are subsequently replaced by obligate anaerobes. The latter support the optimal fermentation of solid food [7, 8].

The abiotic BE together with its indoor microbiome (reviewed by Chase et al., 2016; Karen C. Dannemiller, 2019; Gilbert & Stephens, 2018; Horve et al., 2020; Kelley & Gilbert, 2013) is considered to have an important role in the development and maintenance of a healthy human microbiome [1]. Populations in urban areas of high-income countries currently spend on average 90% of the day indoors; therefore, the abiotic BE and its indoor microbiome represent one of the major reservoirs of environmental microbes [14]. This reservoir plays a vital role in feeding and shaping the human microbiome early in life (see A. W. Brooks et al., 2018; B. Brooks et al., 2014), but it also likely contributes to recolonisation after disease and infection or antibiotic treatment [17, 18].

Cross-talk between the human being and the BE can have both positive and negative aspects. On the one hand, indoor microbes could trigger immediate health issues under rare circumstances. This aspect is of particular concern in the course of pathogen and resistance transmission in hospital environments [19–21]. On the other hand, several studies have pointed out the beneficial effects of a healthy indoor microbiome and presented the observation that the presence of animal-associated and farm-like indoor microorganisms in rural areas is linked to decreased asthma and allergy risk [22–24].

For microbes to be transmitted successfully between hosts via the built environment, they have to be able to survive outside of their natural habitat (“ex-host” survivability) [1, 25]. The majority of human commensals are subject to a number of different stressors, including UV radiation, adverse temperature and desiccation [1]. For gut commensals, the exposure to atmospheric oxygen is a critical factor, as an estimated 99% of all microorganisms that thrive in the distal colon are obligate anaerobes [26]. The extent of aero-tolerance among species varies, and strictly anaerobic taxa are usually unable to actively replicate outside of the gastrointestinal tract (GIT; Ley et al., 2008; Meehan & Beiko, 2014). To overcome this limitation, numerous (anaerobic) bacteria protect themselves through sporulation, as this provides them with the opportunity to escape difficult environmental situations for an extended period of time. It is estimated that about 30% of the genera in the total intestinal microbiota are capable of spore-formation, including *Lachnospiraceae*, *Ruminococcaceae*, *Clostridiaceae* and *Peptostreptococcaceae* [29]. After being transferred to their new environment, spores react to trigger signals (such as bile acids) by germinating, spreading and colonising in the new host thereafter [29].

Non-spore formers, however, must have other strategies that allow them to tolerate environmental stress factors. Strictly anaerobic microorganisms such as *Roseburia* usually die within few minutes of exposure to atmospheric oxygen, as they have not evolved mechanisms that enable them to avoid and repair the damage caused by reactive oxygen species (ROS); i.e. they lack catalases, peroxidases, or superoxide dismutases [30, 31]. Thus, the survival strategies and transmission routes of most of the strictly anaerobic microbiome members, including methanogenic archaea, are largely unknown.

Methanogenic archaea (“methanogens”) represent important components of the human GIT, along with bacteria, viruses and small eukaryotes [32]. These archaea are obligate anaerobic microorganisms and lack the ability to form spores. Despite their oxygen sensitivity, the most common representative, *Methanobrevibacter smithii*, can be detected in almost 96% of the adult population [33]. Interestingly, *M. smithii* is rarely found in younger cohorts [34], and especially uncommon in children born via Caesarean section [35].

In this communication, we address a major question: Can the BE be a potential reservoir for anaerobic, commensal microorganisms? To answer this question, we analysed the indoor microbiome of ten households and compared the microbiome data with the human microbiome data. As the bathroom represents a rich source of GIT-derived microorganisms [36–38], we sampled surfaces near toilets in different households. One major shortcoming of previous studies was that no distinction was made between viable microorganisms (i.e. those with the potential to colonise a new host) and non-viable microorganisms [1]. We addressed this knowledge gap by combining a propidium monoazide (PMA) treatment with high-throughput 16S rRNA gene sequencing. This procedure enabled us to identify the signatures of intact, and thus probably living, microorganisms. As archaeal cells were frequently detected amongst the surviving proportion of the BE microbiome, we analysed the BE archaeome in more detail. The results of this analysis show that human-associated methanogens can survive air exposure and, therefore, could be transferred to another human microbiome through the BE.

## Material and methods

### Sample collection and DNA extraction

Samples were collected from ten different family houses in the vicinity of Graz, Austria (in March 2017). All houses were occupied by at least one adult and five of the houses by families with children. The sampling surface was selected by considering the richest source of GIT-derived microorganisms [36], namely the bathroom. Per household, two areas of 30 cm^2^ in the proximity of the toilet (example shown in Supplementary Fig. 1a) were cleaned with bleach and sterile water and left uncovered and untouched for 7 days. After 7 days, the areas were sampled using a sterile nylon swab (FLOQSwabsTM, Copan, Brescia, Italy) that has been dipped in 0.9% saline (NaCl) solution. Each area was sampled with the swab three times, by rotating the swab every time before sampling the area again (Supplementary Fig. 1b). Metadata collected from the households can be found in Supplementary Table 1. One swab was opened, exposed to the air in an indoor environment and used as a negative control. Additionally, extraction blank controls were added during the DNA extraction step and PCR to identify possible microbial contaminants in the reagents.

The samples were transported back to the laboratory on ice packs on the day of the sampling. One of the two samples from each house, which were collected in parallel, was treated with PMA (Biotium, Inc., Hayward, CA) based on manufactureŕs recommendations and as previously described [39] to differentiate between cells with an intact membrane. Briefly, swabs were transferred to DNA-free 0.9% NaCl-solution and PMA was added to a final concentration of 50 µM and incubated for 5 min on a shaker in the dark. Photoactivation of PMA was performed for 15 min using the PMA-lite™ LED Photolysis device including an intermediate mixing step. PMA is a photo-reactive DNA binding dye which intercalates with free DNA inhibiting downstream DNA amplification thereof. As PMA is not permeable for intact cell walls, a molecular distinction between PMA-masked free DNA from disrupted, most probably dead cells, and DNA from intact, most probably living cells can be made [40]. In the following, we use the term “PMA” to indicate samples treated with PMA (= intact cells only), and the term “non-PMA” for samples not treated with PMA (= both, intact and dead cells/free DNA).

For genomic DNA extraction, the indoor samples were processed using the FastDNA Spin Kit (MP Biomedicals, Germany) according to manufacturer’s instructions. The genomic DNA was eluted in 100 µL elution buffer, and the concentration was measured using Qubit^TM^ dsDNA HS Assay Kit (ThermoFisher Scientific).

### PCR and qPCR

The obtained genomic DNA was used to amplify the V4 region of the 16S rRNA gene using Illumina-tagged primers 515F and 806R (Supplementary Tab. 2). In order to identify the archaeal communities present in the samples, a nested PCR was performed using the primer combination 344F-1041R/Illu519F-Illu806R as described previously [41]. The PCR reaction was performed in a final volume of 25 µL containing: TAKARA Ex Taq® buffer with MgCl2 (10 X; Takara Bio Inc., Tokyo, Japan), primers 200 nM, dNTP mix 200 µM, TAKARA Ex Taq® Polymerase 0.5 U, water (Lichrosolv®; Merck, Darmstadt, Germany), and DNA template (1-2 µL of genomic DNA). The PCR amplification conditions are listed in Supplementary Tab. 3.

The bacterial and archaeal 16S rRNA gene copies were determined using SYBR based quantitative PCR (qPCR) with the primer pair Bac331F-Bac797R and A806f-A958r, respectively (Supplementary Tab. 2). The reaction mix contained: 1x SsoAdvanced™ Universal SYBR® Green Supermix (Bio-Rad, Hercules, USA), 300 nM of forward and reverse primer, 1 µL gDNA template, PCR grade water (Lichrosolv®; Merck, Darmstadt, Germany). The qPCR was performed in triplicates using the CFX96 Touch™ Real-Time PCR Detection System (Bio-Rad, Hercules, USA). The qPCR conditions are given in Supplementary Tab. 3. Crossing point (Cp) values were determined using the regression method within the Bio-Rad CFX Manager software version 3.1. Absolute copy numbers of bacterial 16S rRNA genes were calculated using the Cp values and the reaction efficiencies based on standard curves obtained from defined DNA samples from *Escherichia coli* and *Nitrososphaera viennensis* [42]. The qPCR efficiency was between 85-105%, and the R^2^ values were always above 0.9. Detection limits were defined based on the average Cp values of non-template controls (triplicates) and the corresponding standard curves of the positive controls.

### NGS-based 16S rRNA gene sequencing

Library preparation and sequencing of the amplicons were carried out at the Core Facility Molecular Biology, Center for Medical Research at the Medical University Graz, Austria. In brief, DNA concentrations were normalised using a SequalPrep™ normalisation plate (Invitrogen), and each sample was indexed with a unique barcode sequence (8 cycles index PCR). After pooling of the indexed samples, a gel cut was carried out to purify the products of the index PCR. Sequencing was performed using the Illumina MiSeq device and MS-102-3003 MiSeq® Reagent Kit v3-600cycles (2×150 cycles). The obtained fastq data is available in the European Nucleotide Archive under the study accession number: PRJEB41618.

The fastq data analysis was performed using QIIME2 [43] as described previously [44]. After quality filtering, the DADA2 algorithm [45] was used to denoise truncated reads and generate amplicon sequence variants (ASVs). Taxonomic classification [46] was based on the SILVA v132 database [47] and the obtained feature table and taxonomy file were used for further analysis (Supplementary Tab. 4). The overlapping features from negative controls (DNA extraction and PCR negative controls) were manually subtracted or removed from both the bacterial and archaeal dataset. The reads classified as chloroplast and mitochondria were also removed.

### Human microbiome data

In order to assess to which extent the microbial community indoors is affected and affects the human microbiome, a representative dataset of the human microbiome from several body sites was collected. We took advantage of several previous in-house projects, covering the microbiome of skin, nasal cavity, saliva, urine, vagina, and stool (for more details, please see: Supplementary Tab. 5). All samples were taken from healthy participants and processed in our lab with methods similarly to the current study: DNA was extracted by a combination of enzymatic and mechanical lysis; amplification of the 16S rRNA gene was done using the “universal” primers based on Caporaso et al. (2011) [48] or the slightly adapted primers according to Walters et al. (2016) [49]. Library preparation, as well as sequencing, were performed by the Core Facility Molecular Biology at the Medical University Graz, Austria. Obtained raw reads were processed in parallel to the indoor microbiome data of the current study (Supplementary Tab. 6 and 9).

### Data analysis and statistics

In general, data analysis of the microbial data is based on the indoor data alone (Supplementary Tab. 4 and 9). Comparative analysis between indoor and human microbiomes are based on the mixed dataset (Supplementary Tab. 6 and 10). Bar charts, box plots, and bubble plots are based on relative abundances and were constructed for both the bacterial and archaeal communities at different taxonomic levels using the *phyloseq* [50] and *ggplot2* package in R [51]. Significant differences between microbial taxa were calculated in R using Wilcoxon signed-rank tests for dependent samples (e.g. PMA and non-PMA) or Mann– Whitney *U* test for independent samples (e.g. indoor and human). P values were adjusted for multiple testing using the method after Benjamini & Hochberg (1995). The online-tool Calypso [53] was used to calculate alpha diversity metrics, Linear discriminant analysis Effect Size (LEfSe), the bar plots and principal coordinates analysis (PCoA) plots based on Bray-Curtis. Before analysis in Calypso, the data was normalised using total sum normalisation (TSS) combined with square root transformation. For alpha diversity, reads were rarefied to 5,588 and analysed based on Shannon and richness indices. To test for differences in the beta diversity between sample categories, PERMANOVA analysis was performed in QIIME2 based on Bray-Curtis distances. BugBase [54] was used to predict potential phenotypes such as aerobic, anaerobic, facultative anaerobic and stress-tolerant communities which were present in the PMA and non-PMA treated samples. SourceTracker2 analysis [55] was performed with default settings. Sampling depth was set to 1,000 reads for source and sink, respectively.

A phylogenetic tree was constructed to determine if the sequences belonging to methanogens identified in the analysed samples are of human origin. All sequences classified within the genera *Methanobacterium*, *Methanobrevibacter* and *Methanomassiliicoccus* from our analysed samples were used for creating the phylogenetic tree. Additionally, 16S rRNA gene sequences of species of *Methanobacterium*, *Methanobrevibacter* and *Methanomassiliicoccus* previously identified in the human body or the environment were included from NCBI and two other studies [5, 56]. The alignment was performed using the SILVA SINA alignment tool [57]. The sequences were cropped to the same length using BioEdit and used afterwards to construct a tree based on the maximum-likelihood algorithm with a bootstrap value of 1,000 using MEGA7 [58]. The phylogenetic tree was further processed using the online tool iTOL [59].

### Definition of obligate anaerobic taxa

Bacterial and archaeal taxa were annotated as obligate anaerobic on family and genus level and marked with an asterix *. The categorisation is based on the “list of prokaryotes according to their aerotolerant or obligate anaerobic metabolism” V1.3 [60] and literature review.

### Cultivation and oxygen tolerance test

An oxygen sensitivity test was implemented to determine if methanogenic archaeal strains can survive under aerobic conditions for a certain period of time. Three human-associated methanogenic strains (*Methanosphaera stadtmanae* DSM no. 3091, *Methanobrevibacter smithii* DSM no. 2375, and *Methanomassiliicoccus luminyensis* DSM no. 25720) and a newly isolated strain (*Methanobrevibacter* sp., unpublished) have been tested for their ability to survive under aerobic conditions. The human-associated methanogen cultures were obtained from DSMZ and both, *M. stadtmanae* and *M. luminyensis* were grown in MPT1 medium at 37 °C (Mauerhofer et al., unpublished). The new *Methanobrevibacter* strain was isolated in our lab from a fresh stool sample using ATCC medium: 1340 MS medium for methanogens (https://www.atcc.org/). Both *Methanobrevibacter* strains were grown in MS medium at 37°C.

Cultures were used after 3-7 days of growth as follows: 1 mL culture was transferred to sterile 1.5 mL Eppendorf tubes in the anaerobic chamber and centrifuged at 10,000 x g for 10 min, the cell pellet was washed twice either anaerobically for the controls (anaerobically exposed microorganisms) or aerobically for the oxygen exposed microorganisms with 1 mL sterile 1x PBS and centrifuged again at 10,000 x g for 10 min. After removing most of the PBS, the cell pellet was resuspended in the remaining liquid and exposed to aerobic or anaerobic conditions for different time points: 0 h (30 minutes after transfer), 6 h, 24 h, 48 h, and 168 h (7 days). After exposure, the liquid from the tubes was transferred to a sterile growth medium in Hungate tubes in the anaerobic chamber and the cultures were left to grow for 2-3 weeks. Then, the first optical density (OD) measurements (600 nm) and methane measurements were done using a spectrometer and methane sensor (BCP-CH4 sensor, BlueSens). For the methane measurement, 10 mL of gas-phase was taken from the Hungate tubes using an air-tight glass syringe. For each experiment, two sterile unopened Hungate tubes with media served as control, and two inoculated Hungate tubes with 0.5 mL of the culture served as a positive control. In addition, a tube of 1x PBS served as a control for 168h. Each experiment was performed in duplicates. After OD and methane measurements, microscopic observations were performed to determine the shape of the microorganisms growing in the medium. Oxygen saturation of the PBS was monitored in a control experiment using the FireStingO2 optical oxygen meter (PyroScience GmbH) and the oxygen probe OXROB10. Using the identical methodology as described above we were able to confirm an oxygen saturation in the medium of > 80% at any given time point (0 h, 10 min, 20 min, 30 min, 45 min, 1 h, 2 h, 4 h, 6 h and 24 h) for every strain.

## Results

In this section, we present information about the anaerobic microbial communities detected in the bathrooms of the selected family houses. We wanted to determine whether the indoor environment serves as a source of anaerobic, commensal microorganisms, such as methanogens, which could potentially colonise the human body. To achieve this purpose, we collected samples from two surface areas (∼30 cm^2^) in the bathrooms of ten households.

One of the two samples collected from the BE was treated with PMA in order to mask the DNA of disrupted cells [40]. By comparing non-PMA (untreated) and PMA-treated samples, we could distinguish the overall (non-PMA) microbiome from the intact (i.e. probably alive) microbiome (PMA). This comparison enabled us to draw conclusions regarding the survival of strictly anaerobic microorganisms under exposed conditions. These findings were further supported by the results of subsequent experiments.

Although we present data on all detected microbial signatures, we focused specifically on strictly anaerobic taxa. For convenience, we highlight taxa that mainly consist of strict anaerobes with an asterisk *.

### The human microbiome is the predominant source of microbes in the BE

Using a universal 16S rRNA gene sequencing approach, it was possible to obtain overall more than 380,000 sequences (Bacteria: 99.86%, Archaea: 0.14%), corresponding to 3,684 ASVs and an average of 473 ± 110 ASVs per household (285 ± 50 and 245 ± 80 ASVs in non-PMA and PMA samples, respectively; Supplementary Tab. 4). We observed a high amount of heterogeneity amongst households, with only 26 shared ASVs (0.7%) amongst all households, and 351 ASVs (9.5%) that were shared between at least two households (Supplementary Tab. 7). The 26 ASVs present in all samples include typical skin-associated taxa, such as *Staphylococcus* (11 ASVs) and *Corynebacterium* (8 ASVs), but also *Finegoldia\** (6 ASVs) and *Peptoniphilus*\* (1 ASV).

A total of 32 phyla were observed, and most sequences were classified to four phyla: Firmicutes (52.2% non-PMA, 41.2% PMA), Actinobacteria (20.3% non-PMA, 26.0% PMA), Proteobacteria (18.8% non-PMA, 23.3% PMA) and Bacteroidetes (6% non-PMA, 4.3% PMA) (Fig. 1a). The distribution of sequences classified to these phyla did not change significantly upon PMA treatment. The Linear discriminant analysis Effect Size (LEfSe) method and Wilcoxon signed-rank test were performed, revealing significant differences only in four low-abundance phyla, namely, an increase (Actinobacteria) or decrease (Tenericutes, Fusobacteria, Epsilonbacteraeota) in relative abundance upon PMA treatment (*P* < 0.05, Supplementary Fig. 2).

**Fig. 1:**
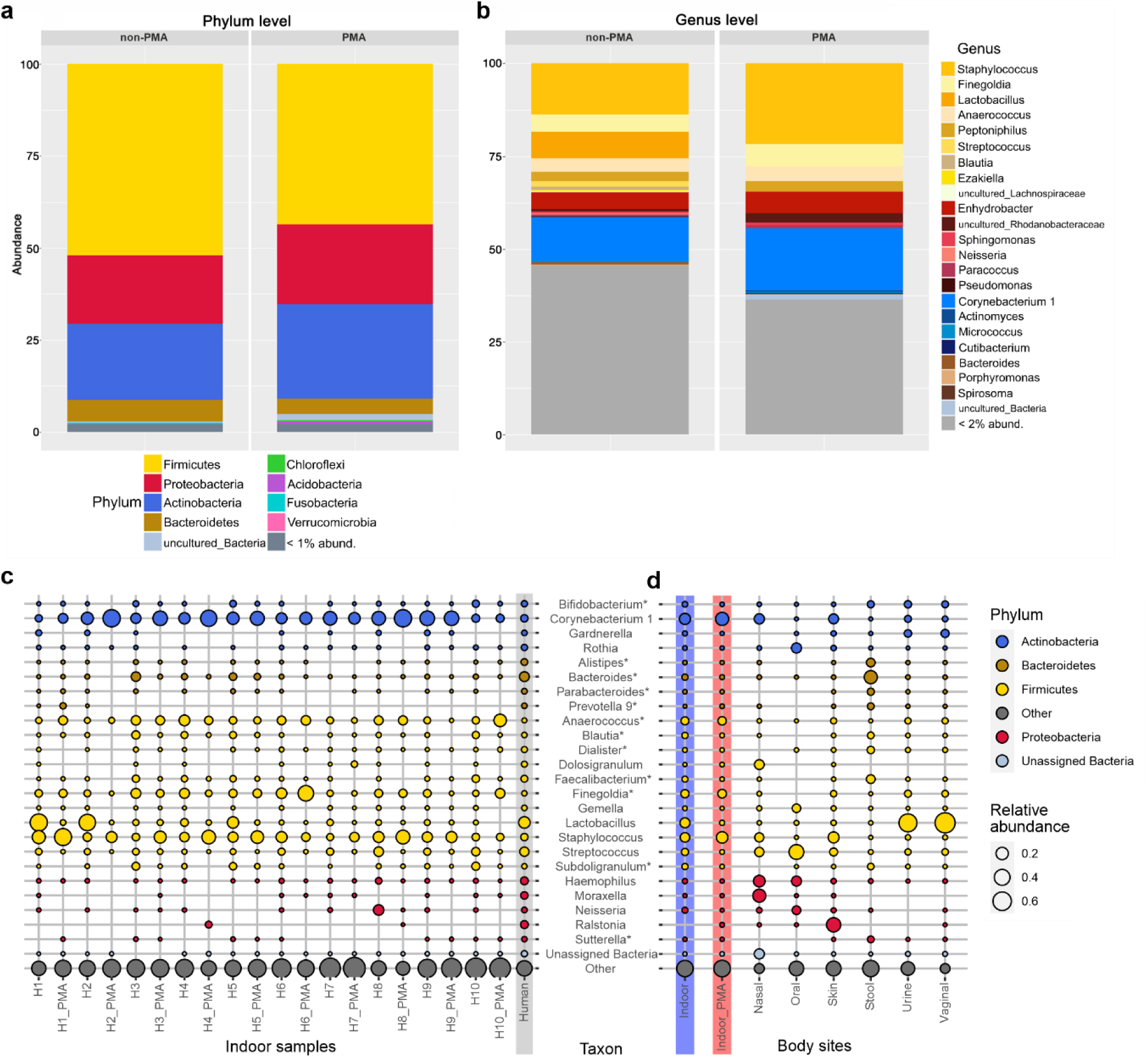
Distribution and relative abundance of bacterial taxa in samples from different households/bathrooms. Bar charts showing the microbial composition of non-PMA- and PMA-treated bathroom floor samples at the **(a)** phylum and **(b)** genus levels. Genera with < 2% rel. abundance are summarised in grey. Bubble plots display the 25 most abundant genera (bubble size reflects the relative abundance): **(c)** Household samples (H1–H10) are depicted individually for non-PMA and PMA treatment together with human samples (grey background) that represent reads from several body sites: nasal cavity, skin, vagina, urine, stool and oral samples. **(d)** Human samples compared to non-PMA (blue background) and PMA (red background) bathroom samples. Genera are coloured according to their taxonomic phylum, and taxa that predominantly contain strict anaerobes are marked by an asterisk.

Genera associated with the human skin (e.g. *Staphylococcus, Corynebacterium, Streptococcus, Neisseria, Micrococcus, Cutibacterium*), gastrointestinal and genitourinary tract (e.g. *Lactobacillus, Finegoldia*, Bacteroides*, Anaerococcus*, Peptoniphilus\**, Lachnospiraceae*) were identified in both non-PMA and PMA samples (Fig. 1c). Notably, *Bifidobacterium*, Rothia, Bacteroides*, Blautia*, Dialister*, Faecalibacterium*, Gemella, Subdoligranulum\** and *Haemophilus* were detected in every single household, but not in all PMA samples, indicating a potential impairment by environmental stressors.

Due to the high proportion of common human-associated commensals, we compared the BE microbiome data collected in this study with human microbiome data that were collected in other studies performed by our lab (see material and methods and Supplementary Tab. 5). The results of a bubble plot analysis of the most abundant genera (Fig. 1d) revealed a specific pattern with respect to the abundance of particular commensals in BE samples or human samples, retrieved from the nasal and oral cavity, skin, stool and vaginal samples. In particular, we observed that signatures of typical human commensals such as *Corynebacterium*, *Anaerococcus*\*, *Dolosigranulum*, *Finegoldia*\*, *Gardnerella* and *Staphylococcus* were highly enriched in non-PMA indoor samples as compared to in human samples (*P*adj < 0.001; Mann– Whitney *U* test), indicating that an efficient transfer to indoor surfaces had occurred.

With respect to alpha diversity, the non-PMA indoor samples showed high Shannon indices, which were comparable to stool samples and even exceeded stool samples in richness by more than two-fold (mean richness is 282 ± 65.3 and 136 ± 30.6 for non-PMA indoor and stool samples, respectively; see Fig. 2). A beta diversity analysis confirmed that all analysed body sites grouped separately (*P*adj < 0.01, PERMANOVA), except for urine and vaginal samples (*P*adj = 0.81; Fig. 2a). However, we did not observe a grouping of indoor samples with a specific human body site, as the clusters differed significantly from one another (*P*adj < 0.01).

**Fig. 2:**
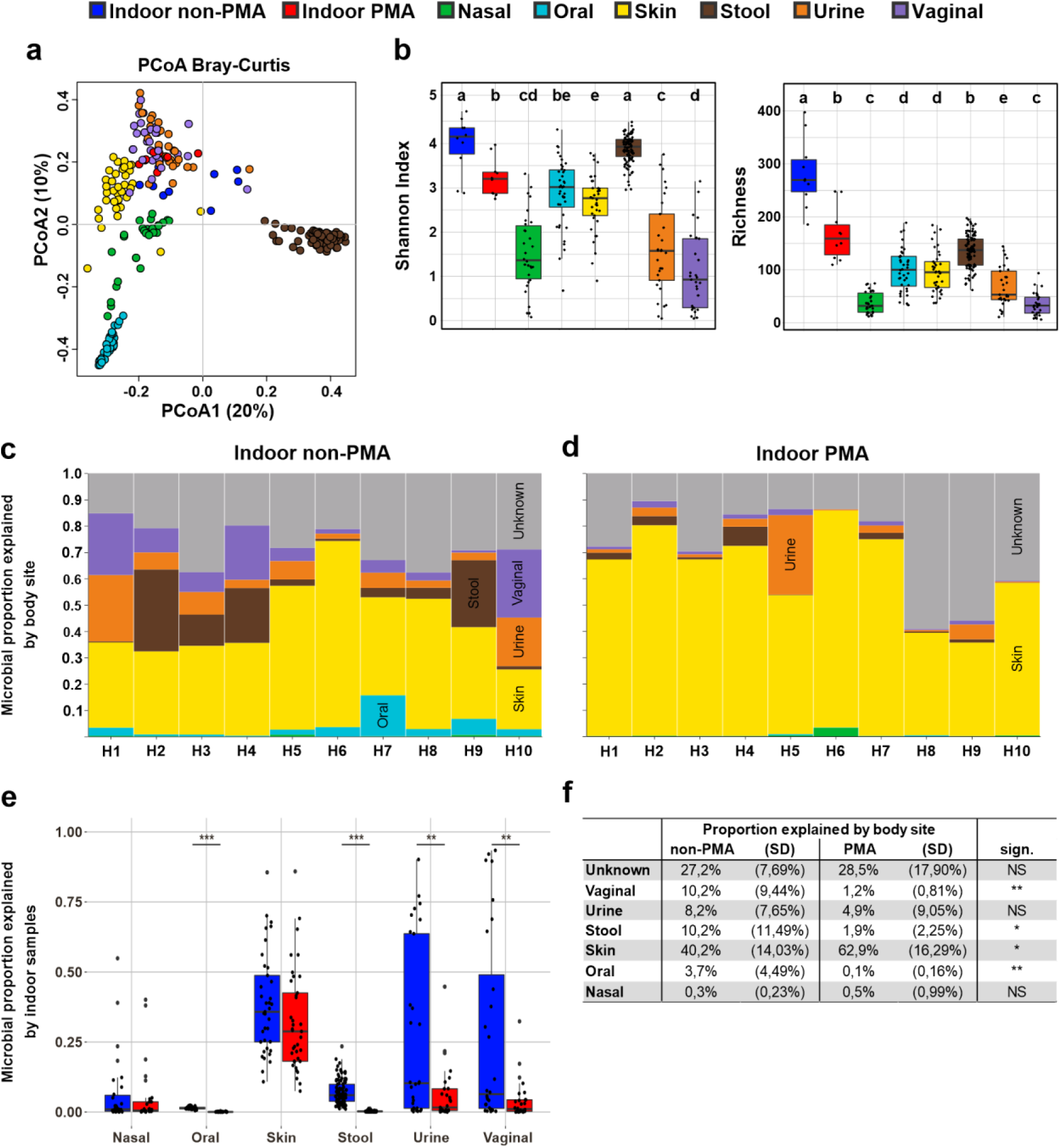
A comparison between the indoor and human microbiome. **(a)** Principal coordinates analysis plots based on Bray-Curtis dissimilarity and **(b)** alpha diversity indices (Shannon, left; richness, right) are depicted for all indoor samples (blue = untreated, red = PMA treated) together with representative human body site samples (green = nasal cavity, light blue = oral, yellow = skin, brown = stool, pink = urine, purple = vaginal). Significant differences are indicated by the different letters above the bars, as defined by a Mann–Whitney *U* test; *P* < 0.05, FDR adjusted (samples that share the same letter do not significantly differ). The proportion of microbes in 10 different bathrooms (H1 – H10) for **(c)** non-PMA and **(d)** PMA samples that can be explained by human body sites or are of unknown origin (grey). Each household is represented by a stacked bar chart, and values were fitted to 100%. **(e)** The proportion of microbes on different body sites that can be potentially explained by the bathroom microbiome; significant differences between treatments were defined by a Kruskal-Wallis test; **, *P* < 0.01; ***, *P* < 0.001. **(f)** The table shows mean values and standard deviations for (d) and (e); significant differences between treatments were defined by performing the Wilcoxon signed-rank test; NS, not significant; *, *P* < 0.05; **, *P* < 0.01.

To determine the level of impact of the human microbiome on our indoor samples, we performed a SourceTracking 2 analysis [55]. Independent of the PMA treatment, about 75% of the indoor microbiome was associated with human sources. These findings indicate that the human body was the dominant source of the microbes detected in bathroom samples (Fig. 3a,b). Nevertheless, depending on the household, we observed large fluctuations in the proportion of human-associated taxa. Especially in PMA samples, this proportion ranges from 41–89%. This may be indicative of a large variation in the frequency and duration of exposure of the tested surfaces to a human being.

**Fig. 3.**
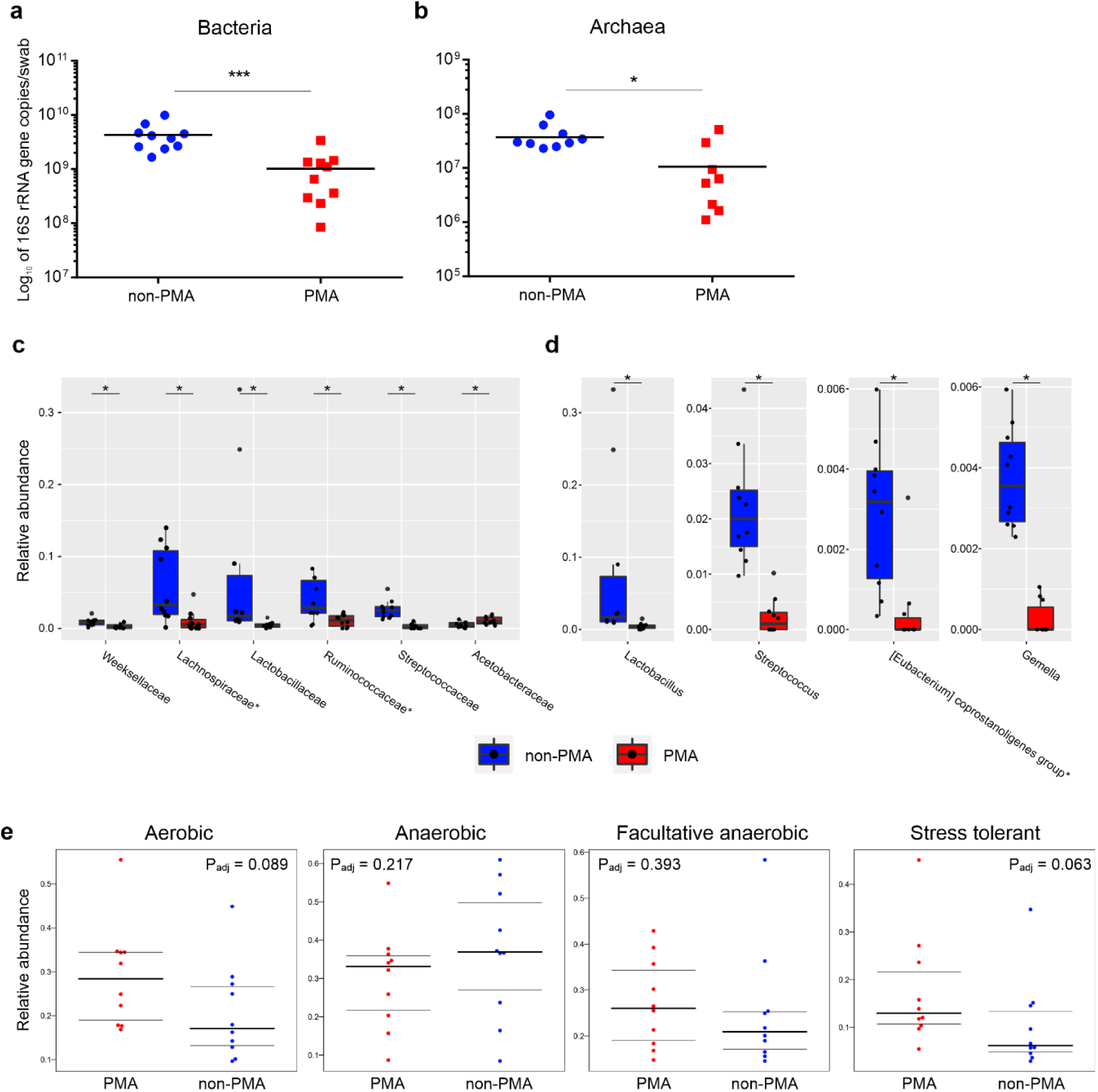
The effects of PMA treatment on prokaryotic abundance and composition. Quantitative PCR analysis of untreated and PMA-treated bathroom floor samples using **(a)** universal and **(b)** archaeal primer pairs. Wilcoxon signed-rank test analysis results, revealing the relative abundance of the 50 and 100 most abundant bacterial **(d)** families and **(e)** genera, respectively, that were significantly affected by PMA treatment. Strict anaerobic taxa are marked by an asterix. Sign.: FDR-adjusted Wilcoxon signed-rank test; *, *P*adj < 0.05; ***, *P*adj < 0.01. **(e)** The relative abundance of different phenotypes as determined through BugBase. Significances were automatically defined by Mann-Whitney *U* test in BugBase.

Upon PMA treatment, we observed a significant reduction in the vaginal, stool, and oral signatures in the surface samples, while the proportion of microbes from human skin significantly increased (*P* < 0.05, Wilcoxon signed-rank test; see Fig. 3d). These results indicate that skin-related microbes possess the greatest ability to survive in the bathroom environment.

*Vice versa*, microbial traces from the indoor environment could potentially serve as a source for the human microbiome as well (Fig. 3c). As has been identified in other studies, we identified that the highest level of exchange occurred between human skin and the indoor and human microbes, whereby the skin shared on average 29% and 32% of the microbial signatures with non-PMA and PMA samples from the bathroom. The overlap with other body sites was generally very low, and especially PMA-treated samples shared on average only about 6% of the taxa with urine, 5% with nasal cavity, 4% with vaginal and < 0.01% with oral and stool samples. Therefore, the impact of the bathroom microbiome on human body sites, with the exception of the skin, seems to be very small or even negligible.

### Oxygen-tolerance primarily determines microbial survival indoors

Human commensals are exposed to several stress factors in the BE; therefore, many of them are perceived as incapable of surviving in this environment over extended periods. We performed PMA treatment to mask DNA from disrupted cells, and in fact, the absolute abundance was significantly affected by PMA treatment (Fig. 3a). Results of the quantitative PCR reveal a 4.2-fold decrease in microbial 16S rRNA gene copy number (Fig. 4c), indicating that less than 25% of all microbial signatures originate from intact, probably living cells. This is also reflected by the significant drop in both of the alpha diversity parameters in PMA samples as compared to those in untreated indoor samples (Shannon: *P*adj = 0.007, richness: *P*adj = 0.0006; Wilcoxon rank-sum tests, Fig. 2b). PCoA plots based on Bray-Curtis distances also showed a significant separation based on the PMA treatment (*P*adj = 0.018) even though indoor samples were more similar to one another than to any human body site (Fig. 2a).

**Fig. 4.**
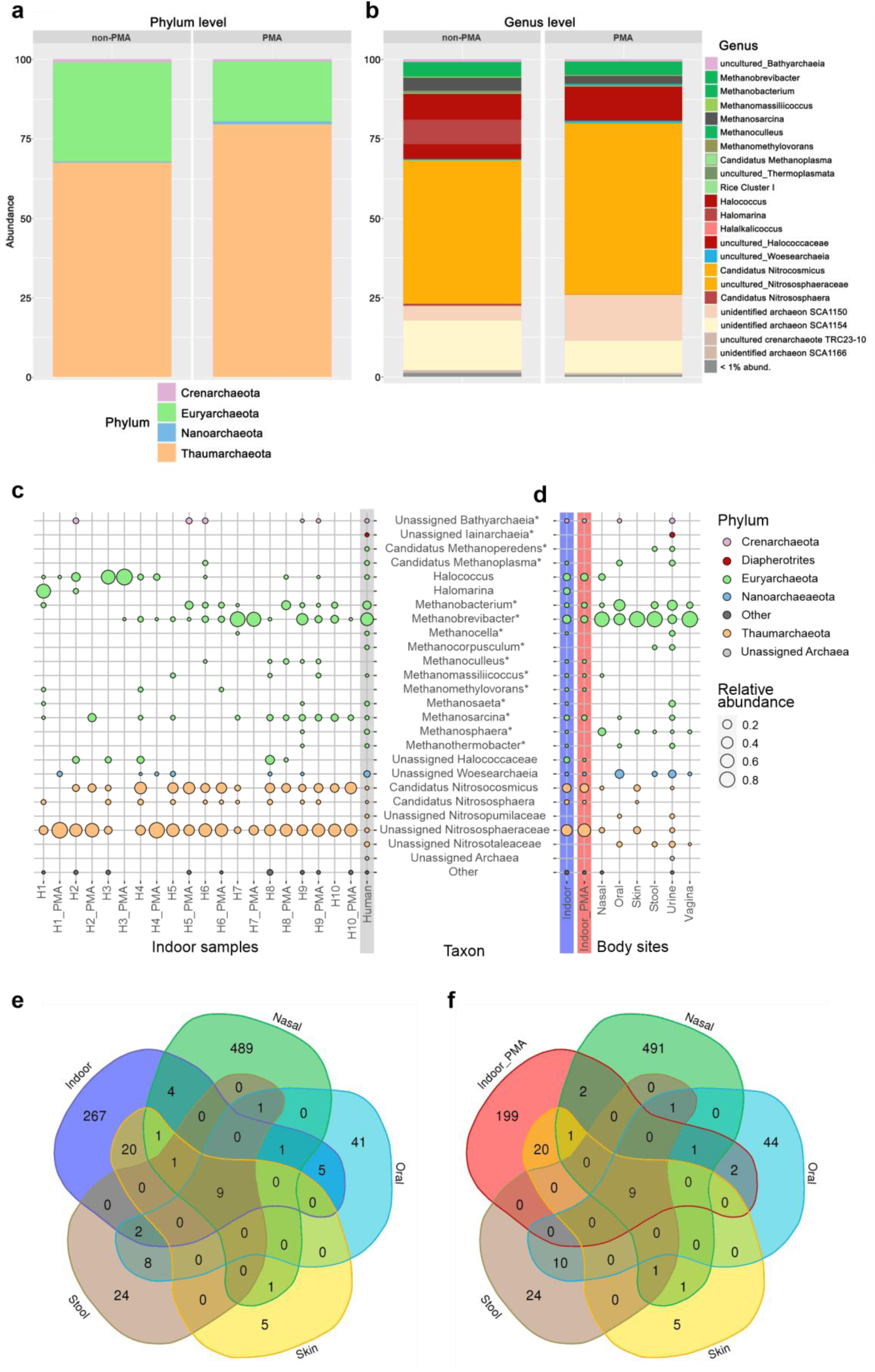
Archaeal community composition in indoor samples. The relative abundance of archaeal taxa among 16S rRNA gene reads from bathroom floor surfaces at the **(a)** phylum and **(b)** genus levels. Bubble plot showing relative abundances of the 25 most-abundant archaeal genera found in bathroom and human samples: **(c)** Household samples (H1-H10) are depicted individually for non-PMA and PMA treatment of human samples (grey background) that represent reads from several body sites including nasal cavity, skin, vagina, urine, stool and oral samples. **(d)** Bubble plots of the 25 most-abundant archaeal genera, displayed according to their original samples. Relative abundance is reflected by the size of the bubbles. Genera are coloured according to their taxonomic phylum, and taxa that predominantly contain strict anaerobes are marked by an asterisk. Venn diagram of shared ASVs in **(e)** non-PMA indoor and **(f)** PMA indoor samples compared to nasal, oral, skin and stool samples (body sites that show the highest numbers of shared ASVs).

LEfSe was performed on PMA and non-PMA samples to determine which taxa are indicative for PMA samples and, therefore, likely to survive in the indoor environment (Supplementary Fig. 3). Representative taxa for PMA were found to be predominantly environmental and aerobic (or aerotolerant) groups, such as *Acidobacteria*, *Micromonosporales*, *Solirubrobacterales*, *Rhizobiales*, *Nocardiaceae*, *Bacillus* and *Sphingomonas,* but also included human skin commensals, such as *Corynebacterium*_1, *Staphylococcus*, or *Rombustia*\*. On the other hand, representative taxa for non-PMA samples were more diverse in terms of their physiological traits and origins. They predominantly included (facultatively) anaerobic, non-spore forming, human-associated groups, such as *Bacteroidales*\*, *Lactobacillaceae*, multiple genera in the families *Lachnospiraceae*\* and *Ruminococcaceae*\*, *Bifidobacteriaceae*\*, as well as in *Christensenellaceae*\*, *Veillonellaceae*\*, *Haemophilus*, *Neisseria* and the *Eubacterium coprostanoligenes* group*. Interestingly, some opportunistic pathogens (*Atopobium*, *Rothia* and *Campylobacter*) were also characteristic for and significantly enriched in the non-PMA samples (Supplementary Fig. 4), which suggests that these taxa might not survive on BE surfaces over extended periods of time.

After PMA treatment, we observed a significant change in the relative abundance of 53 taxa among the most abundant bacterial genera and families (*P* < 0.05, *n*genus = 100, *n*family = 50, Wilcoxon signed-rank test, Supplementary Tab. 8), although only 10 taxa remained significant after performing an adjustment for multiple testing (*P*adj < 0.05, Fig. 3b,c). In accordance with our LEfSe analysis results, we observed a strong reduction of signals originating from non-spore forming, (facultative) anaerobic human commensals, and particularly for *Lactobacillus*, *Streptococcus*, *Gemella* and the *Eubacterium coprostanoligenes* group*. *Ruminococcaceae* and several genera within this family also showed a strong reduction in relative abundance upon PMA-treatment and seemed to be susceptible to stress factors in the BE, whereas taxa in the family *Acetobacteraceae* and other commensals such as *Bacillus* and *Staphylococcus* appear to be extraordinarily well-suited for survival under BE conditions.

BugBase [54] was used to validate the observed phenotypes for non-PMA and PMA samples (Fig. 3d). As compared with non-PMA samples, the PMA samples showed an increase in the relative abundance of aerobic (*P*adj = 0.089) and facultative anaerobic communities (*P*adj = 0.393), and a slight decrease in the abundance of anaerobic communities (*P*adj = 0.217) (Fig. 3d). Interestingly, the abundance of taxa with stress-tolerant phenotypes increased in the PMA samples (*P*adj = 0.063), indicating that microorganisms that could survive in the indoor environment might be better equipped to deal with BE-associated stress factors.

### Archaea are an integral component of the BE

An archaea-centric approach [41] was taken to explore the archaeal communities present on the sampled surfaces. Overall, we obtained about 376,000 archaeal sequences in both PMA and non-PMA samples, which corresponded to 373 ASVs (on average 70.2 ± 22.8 ASVs/ house; 53.6 ± 16.8 ASVs/ non-PMA samples; 35.9 ± 22.1 ASVs/ PMA samples; Supplementary Tab. 9). The archaeal ASVs were classified into four different phyla, namely, Euryarchaeota (36.6% relative abundance in non-PMA samples, PMA: 35.0%), Thaumarchaeota (non-PMA: 62.2%, PMA: 64.1%), Crenarcheota (non-PMA: 0.8%, PMA: 0.5%) and Nanoarchaeota (non-PMA: 0.4%, PMA: 0.5%) (Fig. 4a).

On a family level, most reads were classified into the *Nitrososphaeraceae* (non-PMA: 62.1%, PMA: 64.1%), which is also the only archaeal family that was found in every household included in this study (Fig. 4c). In the *Nitrososphaeraceae*, the dominant genus identified was *Candidatus Nitrosocosmicus* (non-PMA: 22.5%, PMA: 15.9%), but sequences classified within *Candidatus Nitrososphaera* (non-PMA: 0.9%, PMA: 0.2%) were also identified (Fig. 4b). The second-most abundant family was *Halococcaceae* (non-PMA: 19.7%, PMA: 28.7%), with *Halococcus* (non-PMA: 13.7%, PMA: 28.6%) identified as the predominant genus. One particularly fascinating finding was that oxygen-sensitive methanogenic taxa were detected in every household, but at a low relative abundance (*Methanobacteria*\* non-PMA: 3.7%, PMA: 3.5%; *Methanomicrobia*\* non-PMA: 4.0%, PMA: 2.6%), with *Methanosarcina*\* (non-PMA: 3.2%, PMA: 1.8%), *Methanobrevibacter*\* (non-PMA: 2.4%, PMA: 0.2%), *Methanobacterium*\* (non-PMA: 1.3%, PMA: 3.3%) and *Methanomassiliicoccus*\* (non-PMA: 0.4%, PMA: 0.2%) identified as the most abundant genera.

### Humans are potentially the source of most archaeal taxa in the BE

The archaeal community composition suggests that some of the archaeal signatures identified on the bathroom surface could be of human origin, as *Thaumarchaeota*, *Haloarchaea* and methanogens have been previously associated with the human body [62]. Thus, we analysed the data obtained from these surfaces together with data obtained by applying the same archaea-centric approach to samples collected from the human body, namely, to stool (*n* = 38), urine (*n* = 43), nasal (*n* = 30), oral (*n* = 26), vaginal (*n* = 16) and skin (*n* = 7) samples (see Supplementary Tab. 5 and 9).

We recognised that indoor and human samples can be distinguished by their abundance and prevalence of *Thaumarchaeota* and *Euryarchaeota*, as *Thaumarchaeota* are highly enriched in indoor samples (*P*adj = 3.5*10^-11^; see Supplementary Fig. 5; Mann–Whitney *U* test), and *Euryarchaeota* are highly enriched in human samples (*P*adj = 1.3*10^-5^). This especially applies to signatures from *Candidatus Nitrososphaera* (*P*adj = 2.9*10^-26^) and *Cand. Nitrosocosmicus* (*P*adj = 1.1*10^-24^) that were rarely abundant in samples from the human body, but were often found in the analysed indoor samples. In addition, the relative abundance of unassigned *Nitrososphaeraceae* that were found in skin, urine and nasal samples was significantly higher (*P*adj = 1.0*10^-20^) in indoor samples.

Nevertheless, several archaeal genera were found in both indoor samples and human-associated samples. For example, reads classified as *Methanobacterium*\*, *Methanobrevibacter*\*, *Methanosarcina*\*, *Halococcus*, unassigned *Nitrososphaeraceae* and *Woesearchaeia* were present in both kinds of samples, indicating an overlap and that exchange is occurring between both microbiomes.

Venn diagrams were created to illustrate the number of shared archaeal ASVs between the indoor environments and all analysed body sites. Each sample type contained a certain number of specific ASVs (e.g. nasal samples: 489 ASVs, oral: 41 ASVs, skin: 5 ASVs, stool: 24 ASVs, indoor samples: 267 ASVs). The vaginal samples revealed only a small overlap with other samples with respect to the archaeal ASVs (15 unique ASVs, two shared with urine samples). Notably, nine ASVs (*Methanobrevibacter*\*) were shared across all sites, and the skin samples shared the most (20) ASVs with the indoor environment (four *Candidatus Nitrosocosmicus* ASVs and 16 ASVs uncultured *Nitrososphaeraceae*) (Fig. 4e,f).

In the next step, we examined the signatures of methanogenic archaea in more detail, as we had used them as specific model microorganisms in our study. We were interested in determining whether the methanogens identified in the indoor samples were of human origin or were associated with archaeal signatures from the natural environment. Therefore, we first extracted methanoarchaeal sequences from our indoor datasets (PMA and non-PMA) and used these together with reference sequences from different environments that were available from official databases to generate a phylogenetic tree (Fig. 5).

**Fig. 5.**
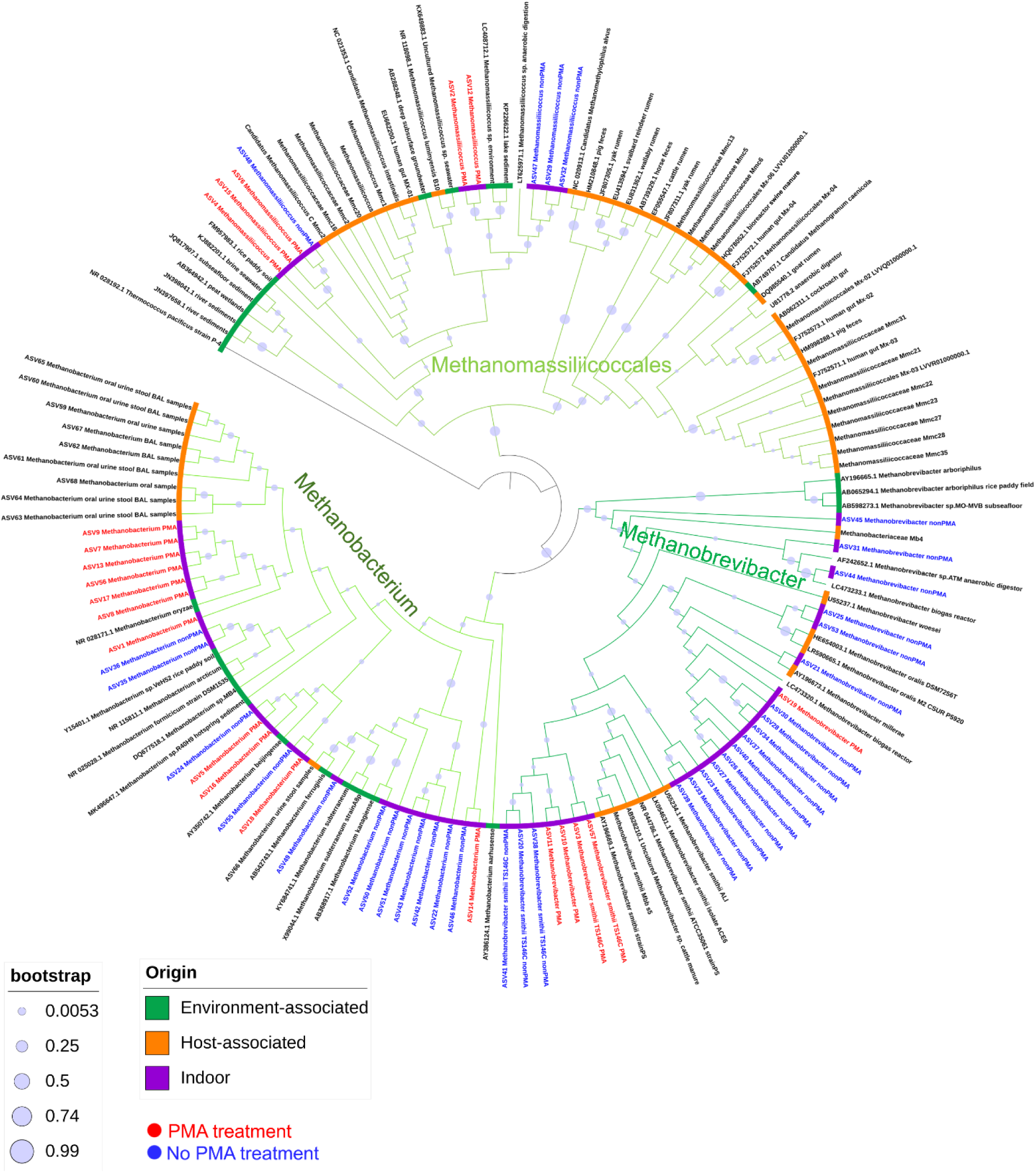
Phylogenetic tree based on the sequences obtained from our study, NCBI and two other studies [5, 56]. The circle indicates the origin of the sequences used to create the tree (see legend). The branches of the tree were coloured in different shades of green according to the taxa they represent.

Most of the *Methanobrevibacter* sequences identified in the indoor samples clustered together with host-associated (human (*M. smithii*\*, *M. oralis*\*), bovine/ovine (*M. millerae*\*), or biogas-reactor/anaerobic-digestor-derived sequences, which are believed to be of holobiont origin. Due to the fact that *Methanobrevibacter*\* is considered to be a strongly host-associated taxon [32], we hypothesise that our detected *Methanobrevibacter*\* signatures are mostly derived from human body-sites and only rarely from other, non-human environments.

Borrel et al. (2017) showed that nearly all human-associated *Methanomassiliicoccales*\* DNA sequences clustered in a host-associated clade together with other *Methanomassiliicoccales*\* sequences identified in animals, with the exception of two taxa, namely *Candidatus Methanomassiliicoccus intestinalis*\* and *Methanomassiliicoccus luminyensis*\*. These two taxa have been identified and isolated from human samples, but they cluster together with sequences of microorganisms that are mainly associated with the soil and sediment environment (“environmental clade”). This clearly shows that some environmental *Methanomassiliicoccales*\* taxa can be transferred from the environment to the human body and can colonise the human gut. The sequences assigned to the order of *Methanomassiliicoccales*\* in our samples clustered only in the environment-associated clade and were closer to the *Methanomassiliicoccales*\* sequences that have been identified in soils, sediments and anaerobic digesters. Therefore, the external environment appears to be the most probable source of the *Methanomassiliicoccales*\* sequences identified in our bathroom floor samples.

Signatures of *Methanobacterium*\* are being more closely associated with human origin [32, 62]. Regarding *Methanobacterium*\*, we included sequences from environment- and human-associated taxa in order to determine whether the *Methanobacterium*\* sequences identified on the bathroom floor were of human origin or not. Most *Methanobacterium*\* sequences identified in our human dataset clustered apart from all other *Methanobacterium*\* sequences, with the exception of one ASV. This ASV clustered together with *Methanobacterium ferruginis*\* and ASV49, which was identified in the non-PMA samples. Seven ASVs from the PMA samples clustered with *Methanobacterium oryzae*\* and shared a node with the phylogenetic group formed by the human-associated ASVs.

The results of phylogenetic and abundance-based analyses indicate that most of the identified methanogenic sequences (and particularly *Methanobrevibacter*\* and *Methanobacterium*\*) in the bathroom floor samples are probably indeed of human origin. This especially applies to the identified sequences in indoor samples.

### Human-associated methanogens can survive oxygen exposure for up to 48 hours

Methanogenic archaea are strict anaerobes and are regarded as highly oxygen-sensitive. Nevertheless, in households with methanogenic taxa, we have frequently observed them in both non-PMA and PMA samples, indicating that these taxa display a certain level of tolerance towards stress factors in the BE. Therefore, we performed an experimental analysis to determine whether methanogenic archaea, serving as models for strict anaerobic gastrointestinal microorganisms, were able to survive under aerial oxygen conditions and, therefore, potentially colonise the human body.

Three methanogen strains previously isolated from the human body, namely, *Methanobrevibacter smithii*\* (DSM no. 2375), *Methanosphaera stadtmanae*\* (DSM no. 3091), *Methanomassiliicoccus luminyensis*\* (DSM no. 25720) and one recently obtained isolate (*Methanobrevibacter sp.*\*; unpublished) from human faeces were exposed to aerobic and anoxic conditions over the period of up to 7 days (0 h, 6 h, 24 h, 48 h and 168 h). After exposure, the cells were transferred to fresh anoxic media and were allowed to grow for 2–3 weeks. Growth was then tested by measuring OD600 and methane production. All four strains could survive in the aerobic environment for at least 6 h (Fig. 6). *Methanomassiliicoccus luminyensis*\* was even able to survive for more than 48 h of exposure to aerobic conditions. No growth of *Methanomassiliicoccus luminyensis*\* was detected after 7 d of cultivation, under either anoxic or oxic conditions, indicating that other negative influences (e.g. starvation) potentially hindered the normal outgrowth.

**Fig. 6.**
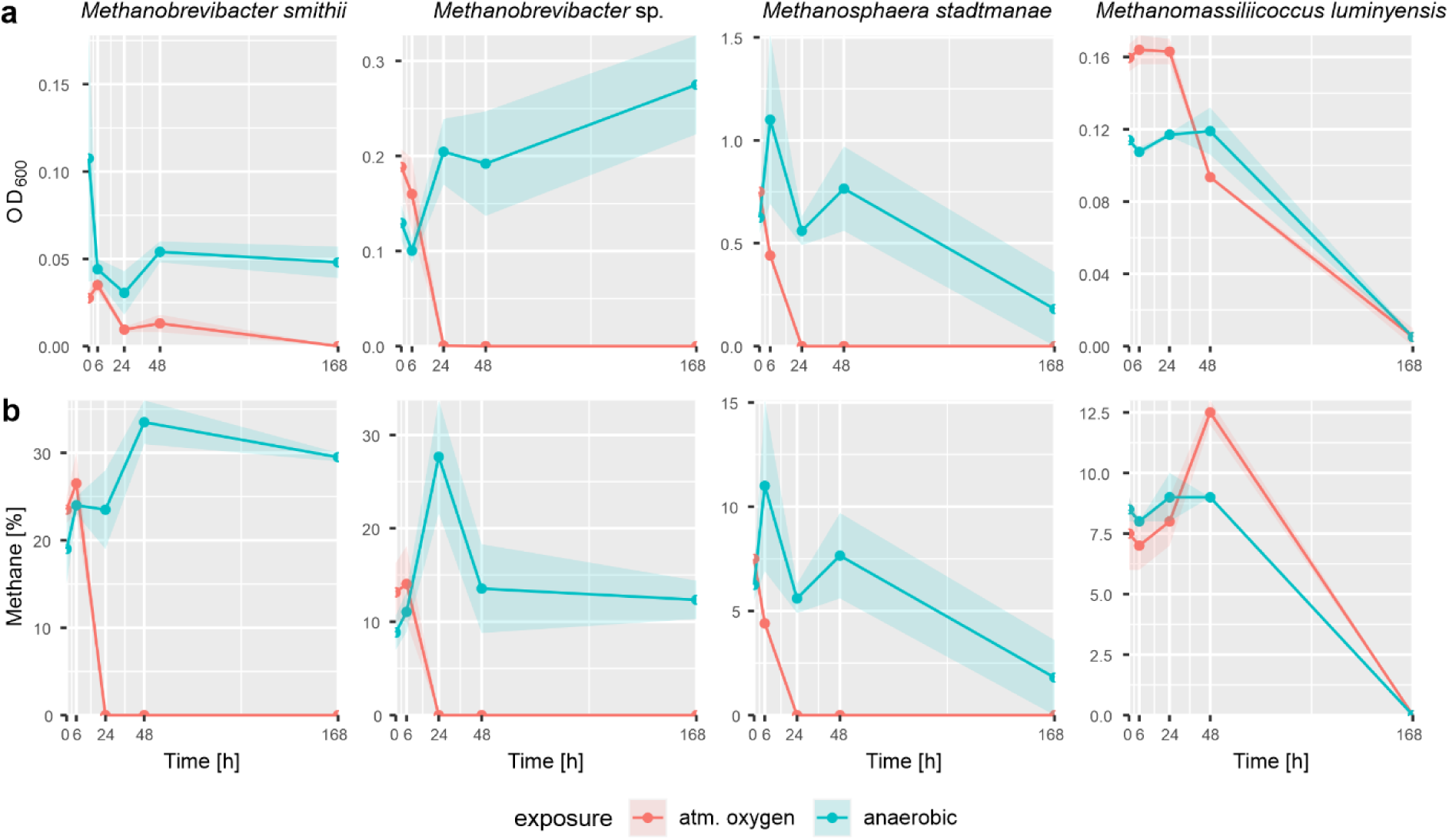
Observed growth of selected human-associated methanogens after exposure to 21% oxygen (air). Plots indicate the growth as determined by **(a)** OD600 measurements and **(b)** methane production of the tested strains after exposure to an aerobic (red) or anaerobic (blue) environment for different periods of time (*n* = 2). The shaded areas represent the standard deviation.

To measure the genomic capacity of methanogens to resist oxygenic stress, we analysed the available representative genomes of *M. stadtmanae*\*, *M. smithii*\*, and *Methanomassiliicoccus*\* (Tab. 1). We specifically searched for keystone genes, as identified in Lyu and Lu (2018), and used the Microbial Genome Annotation and Analysis Platform (MaGe) for detailed comparative genomics [64].

**Tab. 1.**
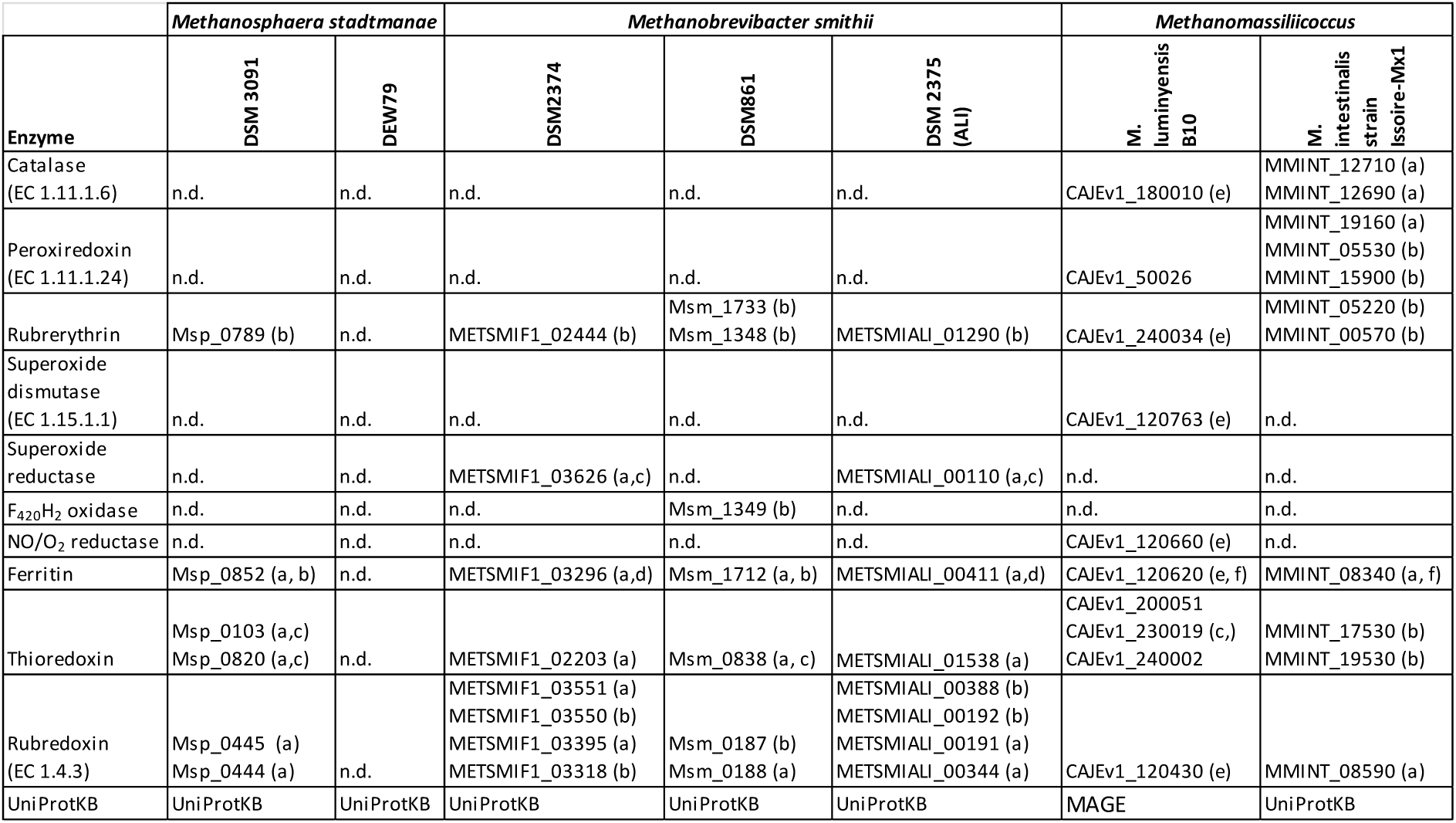
Genes for O2/reactive oxygen species elimination enzymes and their presence in genomes of human-associated methanogens [63]. Explanation of letters in brackets: (a) protein inferred from homology, (b) predicted protein, (c) putative, (d) ferritin-like, (e) function from experimental evidence in other organisms, (f) ferritin, dps family protein. Specific gene identifiers derived from MaGe microscope database [64] and UniProtKB [65] are displayed in each column when present. n.d.: not detected.

All investigated methanogens (except DEW79) revealed the presence of rubrerythrin, ferritin, thioredoxin and rubredoxin in their genomic inventory. The extended survival under oxygenated conditions for *Methanomassiliicoccus*\* was supported by the detection of catalase and peroxiredoxin, as well as superoxide dismutase in the case of *M. luminyensis*\*.

Overall, the experimental and database-derived results show that human-associated archaea have profound capacities to survive under oxygenated conditions for a minimum of 6 hours.

## Discussion

The viability of microorganisms that exist in indoor environments is of great interest [11], as a substantial number of microorganisms are in constant exchange with the human body [15,66– 68]. These interactions between humans and (viable) microbes are arguably of crucial importance for training the immune system and sensitisation in infants [69], as well as the recolonisation of the human microbiome after antibiotic intervention or disease [17, 18].

Our study findings support the hypothesis that the indoor surfaces of normal houses harbour unique microbial communities [66]. The single household profiles are largely dominated by human microbial signatures while environmental taxa play minor roles (for comparison, please see Adams et al., 2015; Chase et al., 2016; Hewitt et al., 2012; Jeon et al., 2013; Rintala et al., 2008), (Fig. 3). This finding is in agreement with the concept of a personalised microbial cloud that mediates the transfer to surfaces [74]. In particular, we identified the skin microbiome as a major contributor to this cloud (∼40%), even in areas close to the toilet, as has been shown in previous studies [37,38,73,75]. Although we observed substantial variation among the microbial communities in the ten households, approximately 30% of the overall microbial community detected on bathroom floors consisted of microbiomes from faecal, vaginal and urinary origins. The presence of those microbial taxa can be explained by the production of aerosols by toilet flushing [76]. However, closing the toilet lid prior to flushing, as is practised in households H4 and H10, did not substantially alter the proportions of contributing body sites. This result raises the question of whether other possible sources of those microbes exist, such as the shower [77] or the sink [36].

Molecular surveys of the BE microbiome usually do not discriminate between viable and dead microbes. As DNA is highly stable in the environment, and microbial signatures remain detectable for a long time, the presence of DNA signatures alone does not serve as a good indicator for viability and metabolic activity per se. Once dispersed, DNA traces of human gut- and skin-associated taxa can still be tracked on public restroom floors and walls after a period of several weeks [37]. Thus, the proportion of relevant microorganisms in the BE might be overestimated when conventional microbiome approaches are taken. Cultivation-based assays, however, are hindered by their technological limitations and might only reflect a non-representative proportion of living indoor microbiota [11].

In this study, we analysed specifically the intact floor microbial community and determined the quality and quantity of the potential microbial ex-host survivors by using PMA treatment, which has rarely been applied to BE environment samples so far [78–80]. One previous specific application was the assessment of living microorganisms in the vicinity of spacecraft, a study that was carried out to understand the risk of contamination for extra-terrestrial research targets (planetary protection; Mahnert et al., 2015; Moissl-Eichinger et al., 2015). Notably, a proportion of 1–45% of the detected microbial community was deemed to be potentially alive under (stringent) cleanroom conditions [79, 81].

Our experiment revealed that the majority of detected microbial signatures originated from disintegrated cells or free DNA. More specifically, we observed a more than 3- and 4-fold decrease in the absolute number of 16S rRNA genes analysed via qPCR for archaea and bacteria, respectively, upon the PMA-treatment of bathroom floor samples (Fig. 3). However, these results also imply that less than 25% of the microbial signatures originate from intact, and likely living, cells that are potentially able to colonise the human body after seven days of regular contact to a human host. The potentially living cells detected in PMA samples could be associated with a predominantly aerobic lifestyle and increased stress tolerance, and these cells could include aerobic, human- and environment-associated, but also spore-forming taxa. The predominance of aerotolerant and spore-forming taxa in PMA samples supports the hypothesis that microbes do not proliferate, but instead persist in the BE as a result of accumulation and dormancy [82].

Whether obligatory anaerobic human commensals can be transmitted between individuals via the BE is still an open question. Notably, microbial taxa such as clostridia might not be directly transmitted from mother to child, even though strains of clostridia are the most abundant bacterial group in the maternal gut [83]. Korpela and de Vos (2018) argued that clostridia may rely on transmission via relatives or non-direct transmission between hosts (i.e. transmission via the environment). This argument is supported by our observation that spore formers, including clostridia, have a perceived increase in survival rate, and seem to be able to survive oxygen exposure and potentially the acidic environment of the stomach.

This kind of horizontal, indirect transfer may be a very rare event in healthy adults, as the transmission success depends on several factors, such as transmission routes, dispersal efficiency, the ability to survive “ex-host”, human colonisation resistance, and the ability of gut commensals to survive gastric and bile acids [1]. These factors are largely unexplored for commensals, but it can be assumed that pathogens and commensals use similar mechanisms. For example, the transmission routes of many pathogens are well-known and primarily include the direct inhalation of aerosols and dust (e.g. *Bacillus anthracis*, *Mycobacterium tuberculosis*) or surface contact (e.g. *Clostridium difficile*\*, *Staphylococcus aureus*, *Enterococcus faecalis*) [11]. It is likely that human commensals also use such mechanisms for transmission. However, such a transmission could not be verified within the scope of our study and has to be evaluated in further surveys by performing a microbial genomic sequence variation analysis to gather evidence for the transfer of commensals between individuals and the BE.

One intriguing results of this study was the detection of archaea as integral components of the BE microbiome. Even though archaea were outnumbered by bacteria nearly 100-fold, they were found in every PMA-treated household sample. Especially taxa of the family *Nitrososphaeraceae* were highly predominant in PMA and non-PMA samples and seem to be relatively stable components of the BE. In contrast, *Euryarchaeota* appeared to be only transient occupants of the indoor microbiome, as many genera were limited to a few households, and their signals frequently disappeared in PMA samples.

An exception to this pattern was observed for the taxon *Methanobrevibacter*\*, which is the most abundant human archaeal genus in the GIT [62]. *Methanobrevibacter*\* signatures were found on all analysed human body sites and in seven out of ten household samples in both PMA and non-PMA samples. These results appear to contradict the general assumption that methanogens are highly oxygen-sensitive [84–87]. However, although taxa of the *Methanobacteriales*\* family seem to be susceptible to ROS [63], analyses of experimental data reveal that methanogens can survive oxygen exposure for periods of several hours to days [88, 89] and are capable to recover from reoccurring aerobic conditions in the environment such as alternated wet and dry rice paddy fields [90–92].

By performing experiments with representative human GIT methanogens, we could confirm that *M. smithii*\* *species* and *M. stadtmanae*\* survived aeration conditions for more than 6 hours, while *M. luminyensis*\* even endured oxygen exposure for more than 48 hours under our experimental conditions (see Fig. 6). The ability of methanogens to endure aerobic conditions was supported by a genomic analysis of common pathways to detoxify ROS. All analysed human-associated methanogens, except *Methanosphaera stadtmanae* strain DEW79, possessed key genes for enzymes such as rubrerythrin, ferritin, thioredoxin and rubredoxin, all of which enable organisms to survive in aerobic environments (see Tab. 1). *Methanomassiliicoccus* sp. further exhibits genes that encode for putative peroxidases and catalases, indicating that this taxon has the potential to deal with ROS, as indicated by our experimental approach.

We are aware, that our study results need to be regarded under the aspect of some methodological limitations. PMA is a powerful tool to mask free DNA but can be affected in its efficacy via physiochemical properties of the sample (e.g. pH, turbidity, optical density) and biochemical characteristics of the microbial community itself (e.g. cell wall structures, natural intake and efflux mechanisms) [40, 80]. In context of this study, we analysed relatively clean, matrix-free BE samples with rather low biomass, suggesting that our results potentially underestimate the amount and diversity of intact microbes in the BE according to a recent evaluation of Wang et al. (2021) [93]. We are thus very confident that the taxa we identified in the PMA dataset originate from intact cells, even though it is still conceivable that some of the taxa found in the PMA dataset are “PMA-resilient”. In our dataset, this may particularly apply for the genera *Corynebacterium* and *Staphylococcus* that have been shown to be fairly unresponsive to PMA treatment [93]. Further limitations include our small sample size and the high level of heterogeneity among the different households which serves as a source of potential bias. In this study, we did not analyse other surfaces/house areas (such as showers or sinks, door handles, dust etc.), which would be of interest in future studies to obtain a better overall picture [36, 73]. Due to the limitations of amplicon sequencing, it was not possible to retrieve functional data for the analyses, i.e. to speculate on the abilities of the microbiome to deal with environmental stress factors. Other techniques, such as shotgun metagenomics, that allow for taxonomic and functional analyses could be employed in future PMA-based BE surveys to obtain such functional data. In addition, environmental, physical parameters such as humidity, temperature, or the frequency of ventilation in the BE should be tracked, as these could help explain microbial survival [94, 95].

By taking into account both 16S rRNA gene amplicon sequencing and experimental approaches, we gathered evidence that supports the hypothesis that methanogens can endure aerobic conditions for a limited amount of time. This introduces a new potential transmission pathway, which is especially interesting with respect to infants, as it presents another way in which they can acquire microbes aside from direct seeding during vaginal birth. It is still not know whether methanogens such as *Methanobrevibacter*\* are acquired perinatal and escape detection early in life or colonise the GIT via another source, such as the BE or dietary products like yoghurt, organic milk and vegetables [96–99]. The latter scenario is plausible, as methanogens co-evolved with animals [56, 100] and, thus, have adapted to the human GIT.

## Conclusions

In conclusion, this study enabled us to use PMA treatment together with molecular quantitative and qualitative methods to successfully assess the survival of Bacteria and Archaea on indoor surfaces. The results indicate that one-fifth of the bathroom surface microbiome is intact or even alive. As even strict anaerobes, such as oxygen-sensitive methanogenic archaea, were found to be intact and potentially alive despite their exposure to the fully aerated environment, we conclude that these microorganisms may represent a valid source for human microbiome constituents, even though a direct colonisation from BE to human needs to be verified in a subsequent study.

## Availability of data and materials

The datasets supporting the conclusions of this article are available in the European Nucleotide Archive (ENA) repository, Primary Accession: PRJEB41618 in https://www.ebi.ac.uk/. Further details can be found in Supplementary Table 4. The R script for the generation of bubble plots was used and adapted from https://github.com/alex-bagnoud/OTU-table-to-bubble-plot.

## Abbreviations

ASV: amplicon sequence variant
BE: built environment
Cp: crossing point
GIT: gastrointestinal tract
LEfSe: Linear discriminant analysis Effect Size
MaGe: Microbial Genome Annotation and Analysis Platform OD: optical density
PCoA: principal coordinates analysis PMA: propidium monoazide
qPCR: quantitative polymerase chain reaction ROS: reactive oxygen species
TSS: total sum normalisation

## Acknowledgements

We thank the participants of our study by providing samples from their bathroom floors. We thank Charlotte Neumann and Christina Kumpitsch for critical proof-reading. Manuela Pausan and Marcus Blohs are students of the local PhD doctoral program “MolMed”. The authors acknowledge the support of the ZMF Galaxy Team: Core Facility Computational Bioanalytics, Medical University of Graz, funded by the Austrian Federal Ministry of Education, Science and Research, Hochschulraum-Strukturmittel 2016 grant as part of BioTechMed Graz.

## Author information

### Authorś contributions

MRP and MB contributed equally to this work. MRP, CME: study design and phylogenetic analysis. MRP: sampling, biological samples processing, collection of metadata, microbiome data analysis, contributed to manuscript writing. MB: analysis of microbiome data and statistical analysis, wrote manuscript. AM: support of microbiome data and statistical analysis. CME: genome mining, wrote manuscript.

## Ethics declarations

### Ethics approval and consent to participate

Ethical approval for the study was obtained from the ethics committee of the Medical University Graz. Written informed consent was obtained from all participants. Ethic approvals for the respective body sites are listed in the Supplementary Table 4.

### Consent for publication

Not applicable.

## Supplementary files

**Supplementary Figure 1.** Sampling area of a representative bathroom floor.

**Supplementary Figure 2.** Analysis of indoor microbiome samples, displaying differences upon PMA treatment on bacterial phyla level. Depicted are taxa that were characteristic to nonPMA and PMA indoor samples. (A) LEfSe analysis, and (B) boxplots of significant different taxa defined by Wilcoxon signed-rank test: *, *P* < 0.05; **, *P* < 0.01.

**Supplementary Figure 3.** LEfSe analysis of bacterial taxa (phylum, class, order, family, genus-level) depicting differences of PMA and non-PMA indoor samples.

**Supplementary Figure 4.** PMA treated indoor samples show a significantly decreased relative abundance for some opportunistic pathogens.

**Supplementary Figure 5.** Comparison of archaeal signatures found on different human body sites (blue) and in the 10 analysed bathrooms (red) on phylum (top) level and for the 25 most abundant genera (bottom). Only samples without PMA treatment were included. Sign.: *, *P* < 0.05; ****, *P* < 0.0001; Wilcoxon signed-rank test.

**Supplementary Table 1.** Metadata collected from all 10 households.

**Supplementary Table 2.** Primer pairs used for archaea and bacteria PCR and qPCR.

**Supplementary Table 3.** The PCR and qPCR conditions for the primer pairs used.

**Supplementary Table 4.** ASV table and data overview for indoor microbiomes.

**Supplementary Table 5.** Summary of additional datasets (human microbiome) used in this study. The study including the saliva samples (bacterial dataset), as well as the study on the nasal cavity (archaeal dataset) have already been published [101, 102].

**Supplementary Table 6.** ASV table of all indoor and human samples, universal dataset.

**Supplementary Table 7.** List of shared taxa among different households.

**Supplementary Table 8.** List of most abundant bacterial genera, families and orders and the respective impact of PMA treatment on their relative abundance, analysed by Wilcoxon signed-rank test.

**Supplementary Table 9.** ASV table of all indoor samples, archaeal dataset.

**Supplementary Table 10.** ASV table of all indoor and human samples, archaeal dataset.

References in Supplementary Tables: [103–108]

