## Supplementary Figures for "The indoor environment - a potential source for intact human-associated anaerobes"

**Supplementary Figure 1:** Sampling area of a representative bathroom floor.

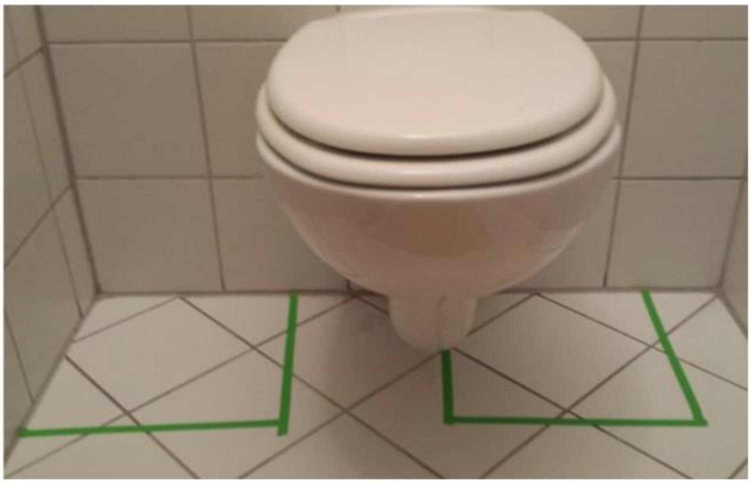

**Supplementary Figure 2:** Analysis of indoor microbiome samples, displaying differences upon PMA treatment on bacterial phyla level. Depicted are taxa that were characteristic to non-PMA and PMA indoor samples. (a) LEfSe analysis, and (b) boxplots of significant different taxa defined by Wilcoxon signed-rank test: *, $P < 0.05$; **, $P < 0.01$.

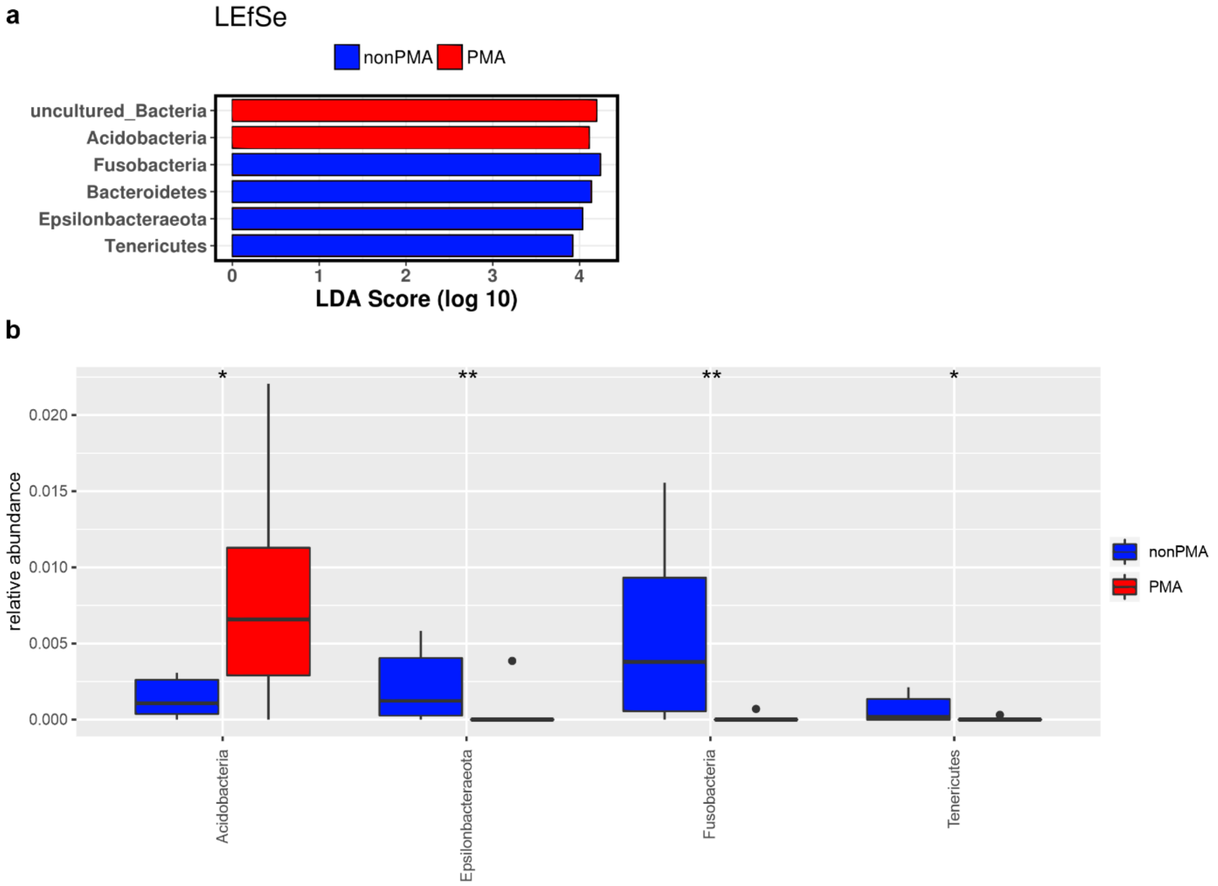

**Supplementary Figures 3.** LEfSe analysis of bacterial taxa (class, order, family, genus-level) depicting differences of PMA and non-PMA indoor samples.

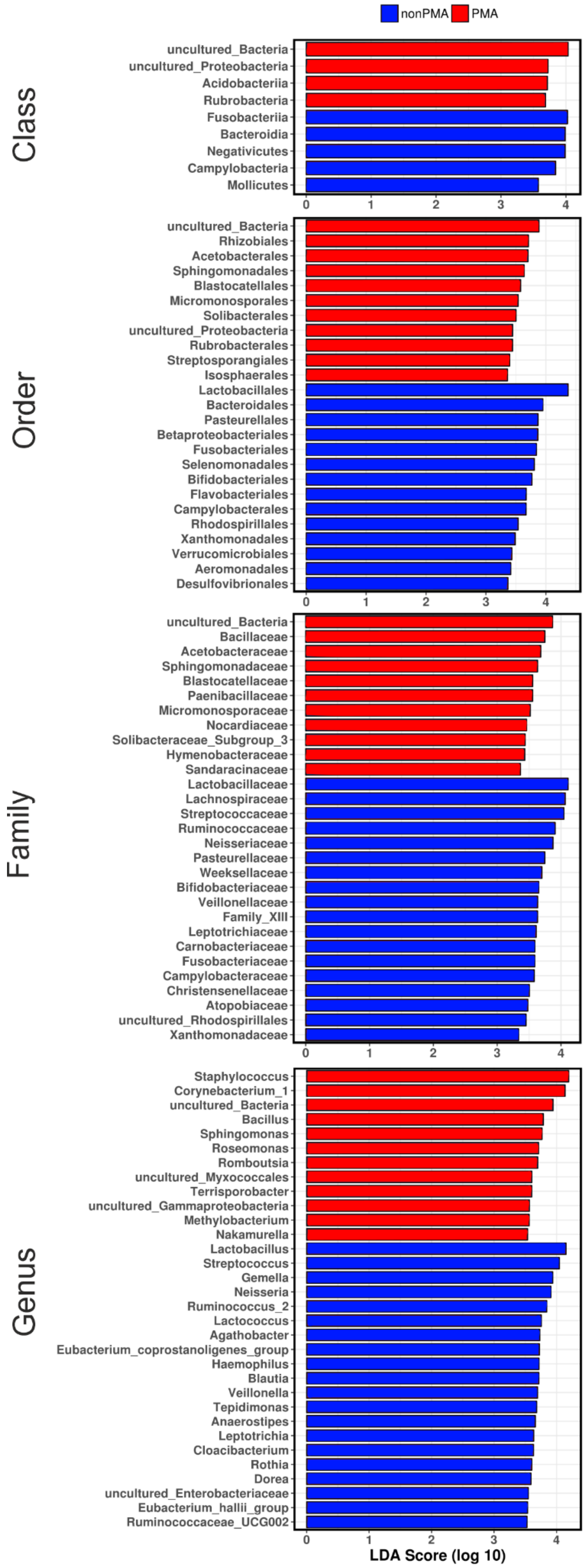

**Supplementary Figure 4**. PMA treated indoor samples show a significantly decreased relative abundance for some opportunistic pathogens.

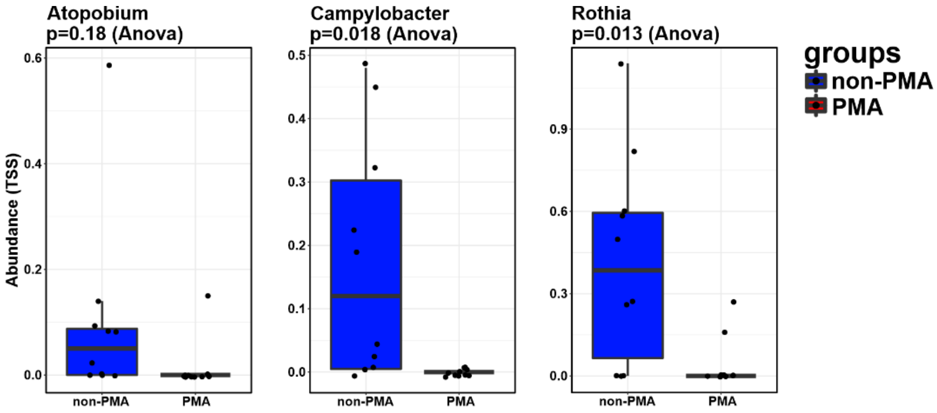

**Supplementary Figure 5**. Comparison of archaeal signatures found on different human body sites (blue) and in the 10 analysed bathrooms (red) on phylum (top) level and for the 25 most abundant genera (bottom). Only samples without PMA treatment were included. Sign.: *, $P < 0.05$; ****, $P < 0.0001$; Wilcoxon signed-rank test.

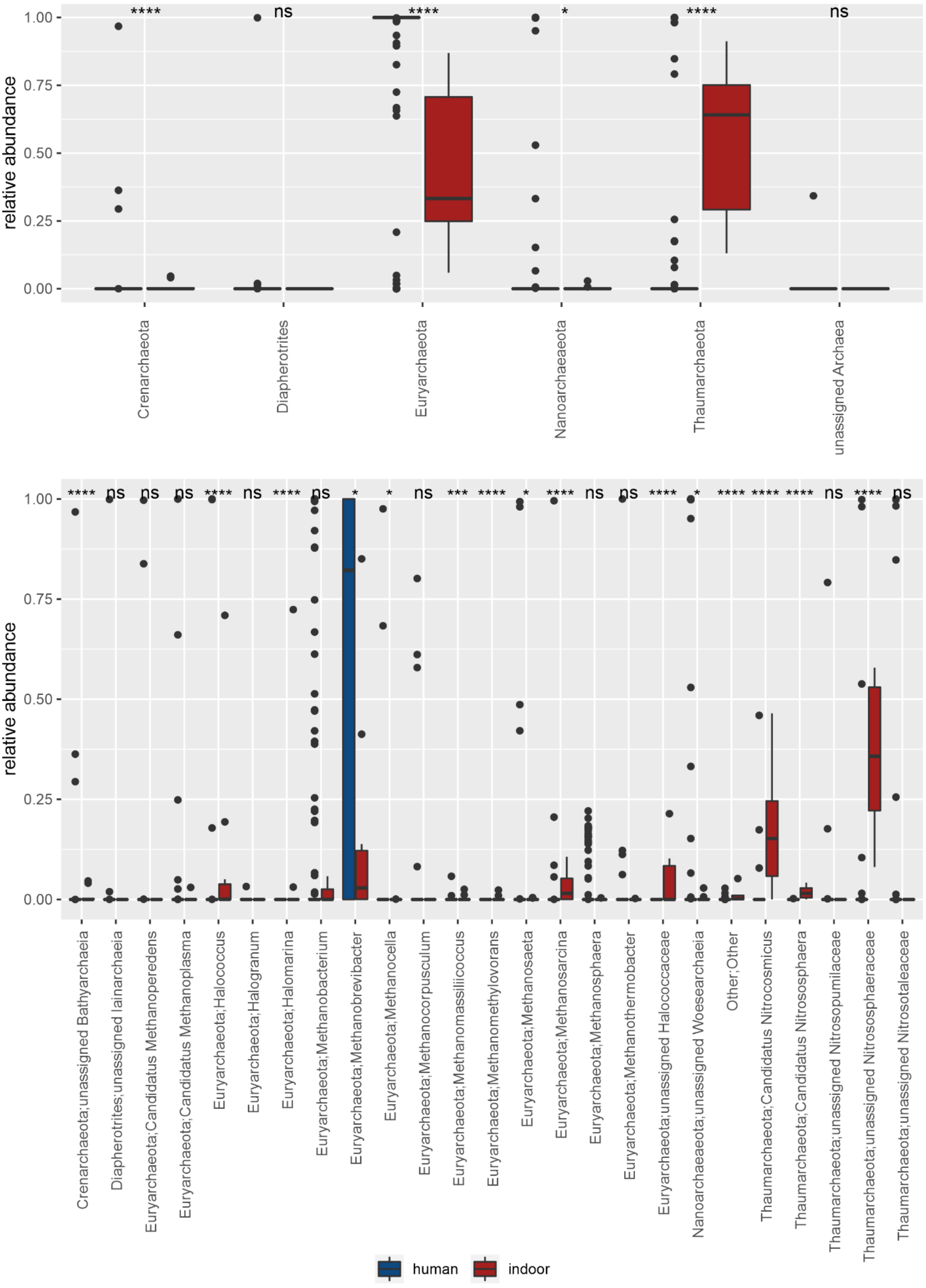
